# Monitoring the eradication of the highly invasive topmouth gudgeon (*Pseudorasbora parva*) using a novel eDNA assay

**DOI:** 10.1101/409821

**Authors:** Chloe Victoria Robinson, Carlos Garcia de Leaniz, Matteo Rolla, Sofia Consuegra

## Abstract

Aquatic Invasive Species (AIS) represent an important threat for Biodiversity and are one of the factors determining the ecological integrity of water bodies under the Water Framework Directive. Eradication is one of the most effective tools for the management of invasive species but has important economic and ecological trade-offs and its success needs to be carefully monitored. We assessed the eradication success of the topmouth gudgeon (*Pseudorasbora parva*), an invasive fish that poses significant risks to endemic aquatic fauna, in four ponds previously treated with the piscicide Rotenone using a novel environmental DNA (eDNA)-qPCR assay. Topmouth gudgeon was detected in all four treated ponds using 750 mL water samples and in three of the ponds using 15 mL samples, despite the eradication treatment. The highly sensitive qPCR assay detected topmouth gudgeon in a significantly greater proportion of sites (77.5%) than eDNA detection methods based on conventional PCR (35%). Our results highlight the difficulties of eradicating invasive fish and the need to incorporate reliable monitoring methods as part of a risk management strategy under the Water Framework Directive.

## Introduction

Invasive species pose one of the main threats to global biodiversity (Sala *et al.,* 2000) through their ability to modify the biological integrity and ecological functioning of native aquatic systems. The presence of Aquatic Invasive Species (AIS) is one of the determinants of the ecological status of European water bodies under the Water Framework Directive (WFD) (Cardoso, Free 2008). Eradication is considered the second most effective tool for the management of invasive species after prevention (Genovesi, Carnevali 2011). However, eradications are typically costly and can have negative impacts on native ecosystems, so the decision to eradicate needs to be taken based on effectiveness, practicality, cost, impact, social acceptability, window of opportunity, and likelihood of re-invasion (Booy et al. 2017). Critical to the application of risk management tools is the ability to detect the target species, even when they are at very low densities (Hulme 2006). Early detection, followed by a rapid response, are critical for the success of eradication programmes (Simberloff, 2014), but these must be monitored to establish their success (Britton et al. 2008). Environmental DNA (eDNA) is increasingly being used for the early detection of invasive and endangered aquatic species (Rees et al., 2014) and is beginning to be used as a monitoring tool under the WFD (Hering *et al.,* 2018).

The topmouth gudgeon (TMG, *Pseudorasbora parva*) is a highly invasive fish originating from Asia that can cause considerable damage to native communities (Britton et al. 2007; Pinder et al. 2005). The species is highly plastic, tolerates a wide range of environmental conditions, and is more fecund than many native fishes in Europe, traits that facilitate its invasion success (Beyer et al. 2007; Britton et al. 2008; Pinder et al. 2005).

In Great Britain, TMG has been reported in 23 locations across England and Wales, the majority being lentic systems, 10 of which are connected to major catchments (Britton et al. 2010; Britton et al. 2007). The species has been classified as highly impactive by the UK Technical Advisory Group on the Water Framework Directive (Panov et al. 2009) due to its rapid reproductive rate, and competition for resources with native fish (Britton et al. 2007). In addition to direct competition, TMG is also a vector for several fish pathogens, including *Sphaerothecum destruens*, the eel nematode parasite *Anguillicola crassu*s, the trematode *Clinostomum complanatum* and the Pike Fry Rhabdovirus (PFR), though the actual impact of these on native fish populations is largely unknown (Britton et al. 2007).

Current methods for managing the spread of TMG have focussed on eradication using the chemical piscicide Rotenone (Allen et al. 2006; Britton et al. 2008), which has been reported to be effective at removing TMG at small spatial scales (Ling 2003). A common problem with the use of piscicides is non-specificity, and the risk of impacting on non-target native fish and invertebrates (Lemmens et al. 2014; Ling 2003). The trade-off between eradication of invasive species and extirpation of native biodiversity needs to be considered, and this can render chemical eradication a non-viable option in some cases (Britton et al. 2011a; Ling 2003).

Invasive species are often difficult to detect using traditional methods both during the early stages of invasion and post-eradication due to their low abundance (Dejean et al. 2012; Jerde et al. 2011; Takahara et al. 2013). TMG undergo boom and bust cycles in novel environments, most likely caused by a combination of biotic and abiotic factors, including temperature fluctuations and predation pressure (Britton et al. 2008; Britton et al. 2007; Copp et al. 2007). Trapping during low abundance or bust cycles may fail to yield any fish and result in false negatives (Davies, Britton 2015). Environmental DNA (eDNA) can be a more effective method for assessing presence/absence, and could inform the success of eradication measures (Davison et al. 2017). eDNA shedding rates (i.e. mucous, faeces, sloughed cells) are higher among fish species (Barnes, Turner 2015) than among other invasive organisms such as invertebrates or amphibians (Ficetola et al. 2008), which facilitates detection and may allow the quantification of fish abundance by quantitative PCR (qPCR), at least in ponds and other closed systems (Klymus et al. 2015; Lacoursière-Roussel et al. 2016).

Rotenone has been used to eradicate TMG from 15 of the 23 TMG confirmed sites in the UK (Brazier 2015), mainly in fishing and recreational lakes, tarns and village ponds across England (Allen et al. 2006; Britton et al. 2008, 2010). In many lakes and ponds with direct connection to a stream screens were installed to prevent dispersal of TMG via rivers before eradication took place (Britton et al. 2008). Subsequent sampling using micromesh seine nets, capable of catching fish >12 mm, failed to detect TMG in all cases since eradication (Britton et al. 2010). One site awaiting eradication in 2014 was the Millennium Coastal Park in Llanelli, Wales, which was reported to have TMG in four ponds within the park (Brazier 2015). Despite two previous eradication attempts at the Millennium Coastal Park during 2012 and 2013, TMG was still detected in one of the main fishing ponds in 2014 (Brazier 2015), possibly due to the presence of a stream which may have allowed recolonization from a nearby pond (Ashpits; (Britton et al. 2009). The use of eDNA was recommended as a more sensitive and robust alternative to netting and trapping to test for the presence/absence of TMG before any further eradication attempts (Davies, Britton 2015).

Previous methods for TMG detection based on eDNA have used end-point (i.e. conventional) PCR (Davison et al. 2017). Here we developed and tested a highly sensitive and specific qPCR high resolution melt curve (qPCR-HRM) assay to test the efficiency of an eradication programme in four ponds and compared the results to traditional end-point PCR. We also assessed the effect of water volume on detection success by using two different protocols involving different water volumes.

## Methods

### Sample sites and eDNA collection

Four ponds (Morolwg, Ashpits, Turbine and Dyfatty) were sampled at the Millennium Coastal Park (Llanelli, Wales) in August 2017. The presence of TMG at Ashpits pond had been confirmed by Rotenone treatment in 2011. Morolwg, Turbine and Dyfatty ponds were also treated with Rotenone in 2012, however due to their proximity to Ashpits pond and the operation of fishery for a local angling group, there were concerns that TMG could have been reintroduced into Morolwg pond. There had been no records of TMG in Dyfatty or Turbine ponds since 2012 (Table 1; Figure 1).

**Table 1.**
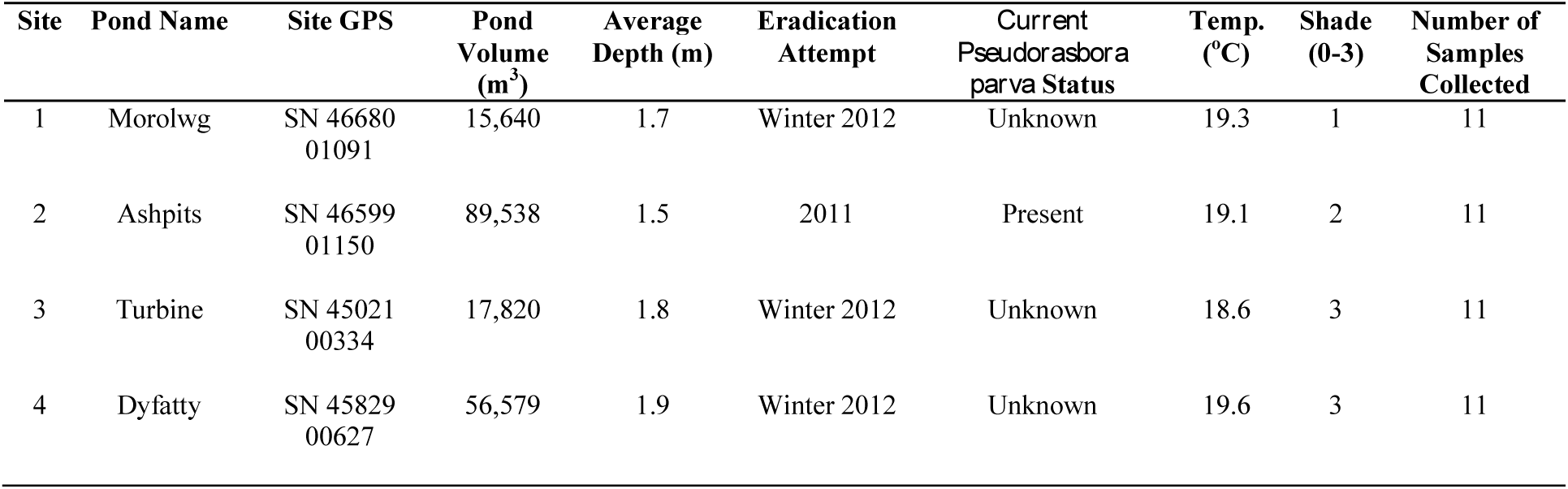
Site information for the four ponds sampled for presence of *Pseudorasbora parva* in South Wales during August 2017

**Figure 1.**
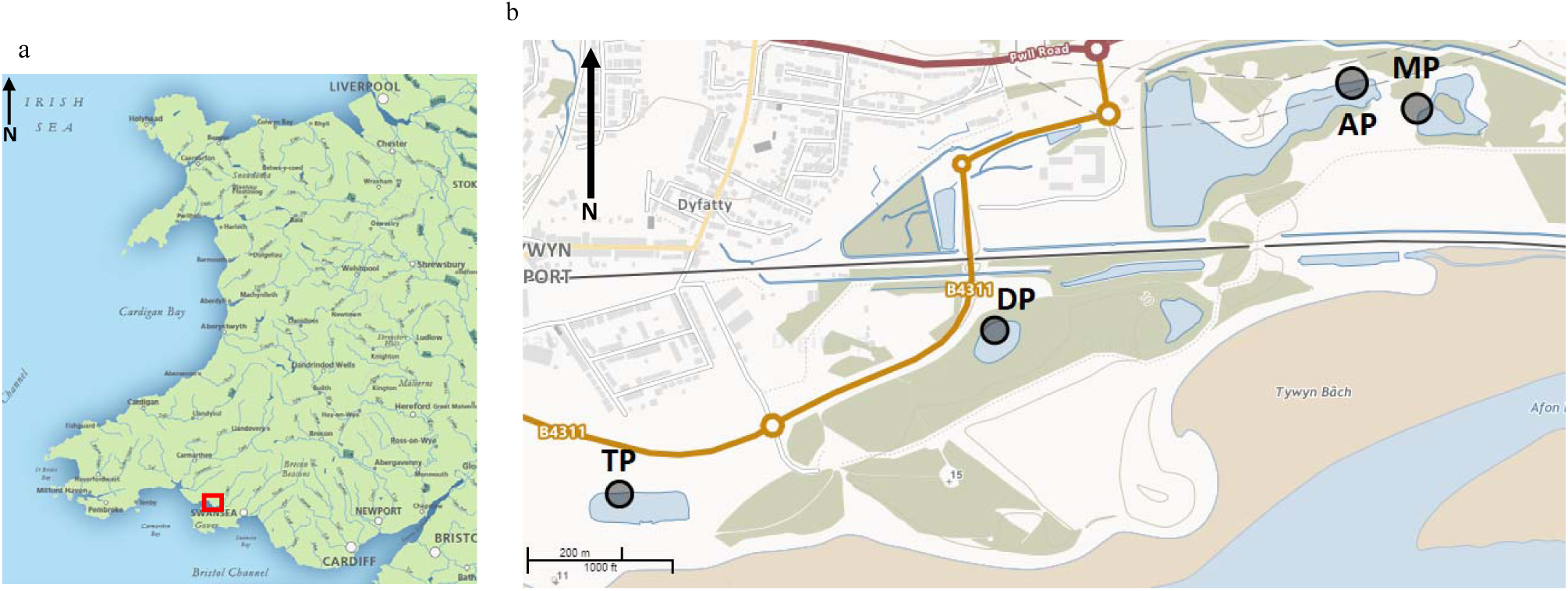
Location of study ponds. A) Location of the Millennium Coastal Park within relation to Wales (red rectangle). B) Four ponds sampled for *Pseudorasbora parva* eDNA Millennium Coastal Park in Llanelli. At each pond (Morolwg Pond, MP; Ashpits Pond, AP; Turbine Pond, TP and Dyfatty Pond, DP) both ten 250 mL water samples were collected in triplicate (750 mL total per sampling point) and one 15 mL water sample were collected at each sampling point.

A sample of 250 mL of pond water was collected in triplicate using a sterile collection ladle before being pooled into a sterile 1L Nalgene bottle (final volume 750 mL) at ten sampling points per pond. Sampling points were separated between 20 and 100 m per pond. An additional water sample of 15 mL was collected at each point per pond as in (Ficetola et al. 2008) to assess the sensitivity of the water volume to detect the target species in ponds. To each 15 mL sample, 33 mL of absolute ethanol and 1.5 mL 3M sodium acetate were added; tubes were kept upright on ice for transportation and subsequently stored at −20 ^°^C until DNA extraction. Ladles were decontaminated by thorough spraying with Virkon^®^ followed by rinsing three times with ultrapure water between study ponds to prevent potential DNA carryover, resulting in false positives. Negative controls were taken at each pond for both methods and before sampling and after decontamination of each ladle before sampling a new pond. Environmental conditions, including water temperature, water depth and extent of cover were recorded at each site (Table 1).

Trapping of TMG was attempted at three time points under permit EP/CW061-E-546/11141/02. In July 2017, five standard minnow traps (20 cm x 20 cm x 60 cm) were 6placed by Natural Resources Wales at Ashpits pond at 1 m depth for seven days, concentrating on the eastern side of the pond where TMG had been previously seen. Traps were baited with fish pellets and checked daily. In February 2018, seine nets and ten standard minnow traps baited with fish pellets and algal-based bait were placed evenly around Ashpits (seine nets and traps) and Dyfatty Ponds (only traps) by Swansea University research staff at a depth of 1 m and checked after 3 hours. The final trapping time point was June 2018 under permit EP/CW061-E-546/12754/01, where ten standard minnow traps were placed around two ponds (Ashpits and Morolwg) over a two-day period, baited with fish pellets and set at a depth of 1 m and checked after 2 hours for Ashpits and after 24 hours for Morolwg. Seine netting was also carried out in Ashpits, Morolwg and Dyfatty ponds. In addition, eight larvae (< 12 mm) were collected by hand-netting from Ashpits (n=5) and Morolwg (n=3) and transported back to the university before being euthanised following Schedule 1 protocol of overdose of 2-Phenoxyethanol.

### Primer design and DNA extraction

Species-specific qPCR primers (PparvaF 5’-CGAGCCCAAATAACAGAGGGT-3’ and PparvaR 5’-CAGGCGAGGCTTATGTTTGC-3’) were designed for TMG using NCBI Primer-BLAST to amplify a 147 bp fragment of the 16S mtDNA gene and checked for cross-amplification using NCBI Primer-BLAST (Ye et al. 2006). Primers were assessed *in vitro* using positive control tissue (caudal muscle) from 15 TMG caught locally in Wales during 2012/13. DNA was extracted using the Qiagen^®^ DNeasy Blood and Tissue Kit (Qiagen, UK), eluted in 200 µl, and amplified in end-point PCR using the following Pparva16S protocol: 95^°^C for 3 min, followed by 40 cycles of 95 ^°^C for 30 s, 61 ^°^C for 30 s and 72 ^°^C for 30 s with a final elongation step of 72 ^°^C for 10 min. All amplified PCR products were checked for the correct amplicon size using a 2% agarose gel electrophoresis.

### qPCR optimisation

Primers were tested *in vitro* for non-specific amplification against closely related species, including common carp (*Cyprinus carpio*), silver rudd (*Scardinius erythrophthalmus*), common bream (*Abramis brama*) and common roach (*Rutilus rutilus*), which are known to inhabit similar water systems (Davison et al. 2017). DNA from these species was amplified in triplicate in qPCR with a positive control consisting of 0.1 ng of topmouth gudgeon DNA (Davison et al. 2017). All non-target species failed to amplify and no subsequent products were produced in HRM analysis. Specific *in vitr°* testing ° f HRM-qPCR was perf° rmed f° r TMG t° c° nfirm the c° nsistency ° f the qPCR pr° duct melting temperature. The annealing temperature ° f *Pparva16S* primers was ° ptimised at 61^°^C and yielded an efficiency of 91.1%, R^2^ = 0.981 (Figure S1; Table S1 and S2). For optimisation, the Pparva16S-qPCR was undertaken using SsoFast™ EvaGreen^®^ Supermix (Bio-Rad, UK) and the cycling protocol began with 15 min of denaturation at 95 ^°^C, followed by 40 cycles of 95 ^°^C for 10 s and 61^°^C for 30 s. A HRM step was applied to the end of RT-qPCR reactions, ranging from 55 ^°^C to 95 ^°^C in 0.1 ^°^C increments to assess the consistency of amplicon melt temperature (tm). The limit of detection (LOD) and limit of quantification (LOQ) were determined through running a dilution series ranging from 5 ng/µl to 5 x 10^−7^ ng/µl, using a TMG DNA pool. HRM analysis was conducted on 15 individuals from three different populations to account for any degree of intraspecific variation in qPCR product melt temperature (tm).

### Analysis of eDNA field samples

Samples of 750 mL of pooled water (3 x 250 mL at each sampling site) from each of the study ponds were filtered the same day of collection through borosilicate glass fibre filters (0.45 µM pore size; Whatman™, Sigma Aldrich, UK) using a vacuum pump (Cole Palmer, UK). DNA filtration took place in a designated eDNA area in a laboratory where no previous TMG DNA or tissue had been handled. Filters from water samples were stored in individual Eppendorfs at −20 ^o^C until subsequent DNA extraction. Filter apparatus was dismantled and decontaminated between samples from different ponds with 10% bleach solution, rinsed with ultrapure water and placed under UV light for 20 minutes. Filtration blanks were obtained by filtering 750 mL of ultrapure water after decontamination of the apparatus to ensure that the cleaning process had been effective. TMG DNA from filter papers was extracted using Qiagen^®^ DNeasy Blood and Tissue Kit (Qiagen, UK), following the Qiagen Blood Spot extraction protocol with an adjustment to the elution volume (from 200 to 50 µl) to maximise DNA yield.

The 15 mL water samples were centrifuged at 6 ^o^C for 45 minutes at 5000g (Ficetola et al. 2008) and the supernatant was poured off to allow DNA extraction from the resulting pellet. DNA pellets were extracted using DNeasy PowerLyzer PowerSoil kit (Qiagen^®^, UK) following a standard protocol with a reduction in elution volume (60 to 50 µl). DNA extractions were undertaken in a dedicated eDNA area within an extraction cabinet equipped with flow-through air system and UV light and to minimise cross-contamination; additionally, a dedicated eDNA laboratory coat and nitrile gloves were worn during the process. All samples were amplified in triplicate in a Bio-Rad CFX96 Touch Real-Time PCR Detection System (Bio-Rad, UK), in 10 µl reactions consisting of 5 µl SsoFast™ EvaGreen^®^ Supermix (Bio-Rad, UK), 0.25 µl each forward and reverse primer, 2.5 µl HPLC water and 2 µl of extracted DNA. Amplifications were carried out in triplicate using the standard Pparva16S-qPCR protocol as described above and only samples which amplified consistently in one of three replicates at the target DNA product tm (78.8^°^C ± 0.3), with a melt rate above 200 −d(RFU)/dT were considered to be a positive result. qPCR reactions were carried out in a separate room from the eDNA extractions room under a PCR hood with laminar flow. We added a TMG positive control to each plate after all the eDNA samples had been loaded and sealed to detect any false positives. As negative amplification controls we used HPLC water which, along with extraction negative controls, were added to the same well location on each plate to test for eDNA contamination. Field samples were also amplified in triplicate in end-point PCR with recently described species-specific primers (Davison et al. 2017), using the *Pparva16S* PCR protocol, and subsequent PCR products were checked for the correct amplicon size using a 2% agarose gel. Any positive reactions for TMG DNA with both primer sets were sequenced on an ABI Prism 377 sequencer to confirm species identity.

### Analysis of larvae

DNA was extracted from a total of eight larvae from Ashpits (n=5) and Morolwg (n=3) ponds using Qiagen^®^ DNeasy Blood and Tissue Kit (Qiagen, UK). Larvae DNA was eluted in 200 µl, and amplified using both end-point PCR and the new qPCR Pparva16S protocol. All amplified end-point PCR products were checked for the correct amplicon size using a 2% agarose gel electrophoresis and were sequenced on an ABI Prism 377 sequencer to confirm species identity.

### Statistical analysis

We employed a generalized linear modelling approach in R v.3.4.3 (R Core Team, 2017) to model detection success (i.e the proportion of sites that tested positive for topmouth gudgeon at each pond) as a function of assay type (three assays: conventional PCR on 750 ml of water, qPCR on 15 mL of water, and qPCR on 750 mL of water) and pond identity (n: 4 ponds). We considered that topmouth gudgeon was present at a site if one of the three replicates tested positive for that site. A quasibinomial log-link was used to correct for overdispersion.

## Results

### Sensitivity and detection limits

No TMG were caught in any of the two trappings at Ashpits pond during July 2017 and February 2018. The limit of detection (LOD) for TMG DNA was 0.005 ng/µl, determined through a 10-fold dilution series in qPCR. The detection threshold for positive control DNA was between 26 and 38 cycles (Table S1) and the product melt temperature was consistent across the entire dilution range (Table S2). In comparison, the primer sensitivity of end-point PCR with species-specific COI primers (((Davison et al. 2017) at 40 cycles) and the newly designed *Pparva16S* primers amplified TMG DNA pools at 0.05 ng/µl and above, highlighting the greater sensitivity of qPCR, with *Pparva16S* primers producing a higher yield of DNA based on band brightness (Figure S2).

### Detection success

Larval DNA sequencing and qPCR profiles confirmed that two larvae caught from Ashpits pond matched 100% with TMG on BLAST, despite lack of adult TMG being caught in traps and seine netting at trapping events. No positive amplification was obtained for any of the other larvae. Results of qPCR confirmed the presence of TMG DNA in all ponds (Figure S3; Table S3). Morolwg Pond (MP) yielded the highest proportion of sampling sites amplifying for TMG (8 out of 10 sites), whereas Turbine Pond (TP) had the lowest proportion of positive sampling sites (6/10; Table 2). The 15 mL water samples successfully detected TMG in three out of the four ponds, with no TMG DNA being detected in Dyfatty Pond (Figure S4; Table S4). In comparison, results of end-point PCR with species-specific COI primers (Davison et al. 2017) failed to produce any positive amplification for TMG unless the PCR reaction was undertaken with 40 cycles. End-point PCR results also showed that TMG DNA was present in all four ponds, however most positive samples were only observed in one of the three triplicates and product bands were faint (Table 2; Figure 2). Sequencing of the 350bp (Davison et al. 2017) and 147bp products produced a 100% species match on BLAST (Ye et al. 2006), with target species (for both primer sets), confirming species presence in all positive amplifications.

**Table 2.**
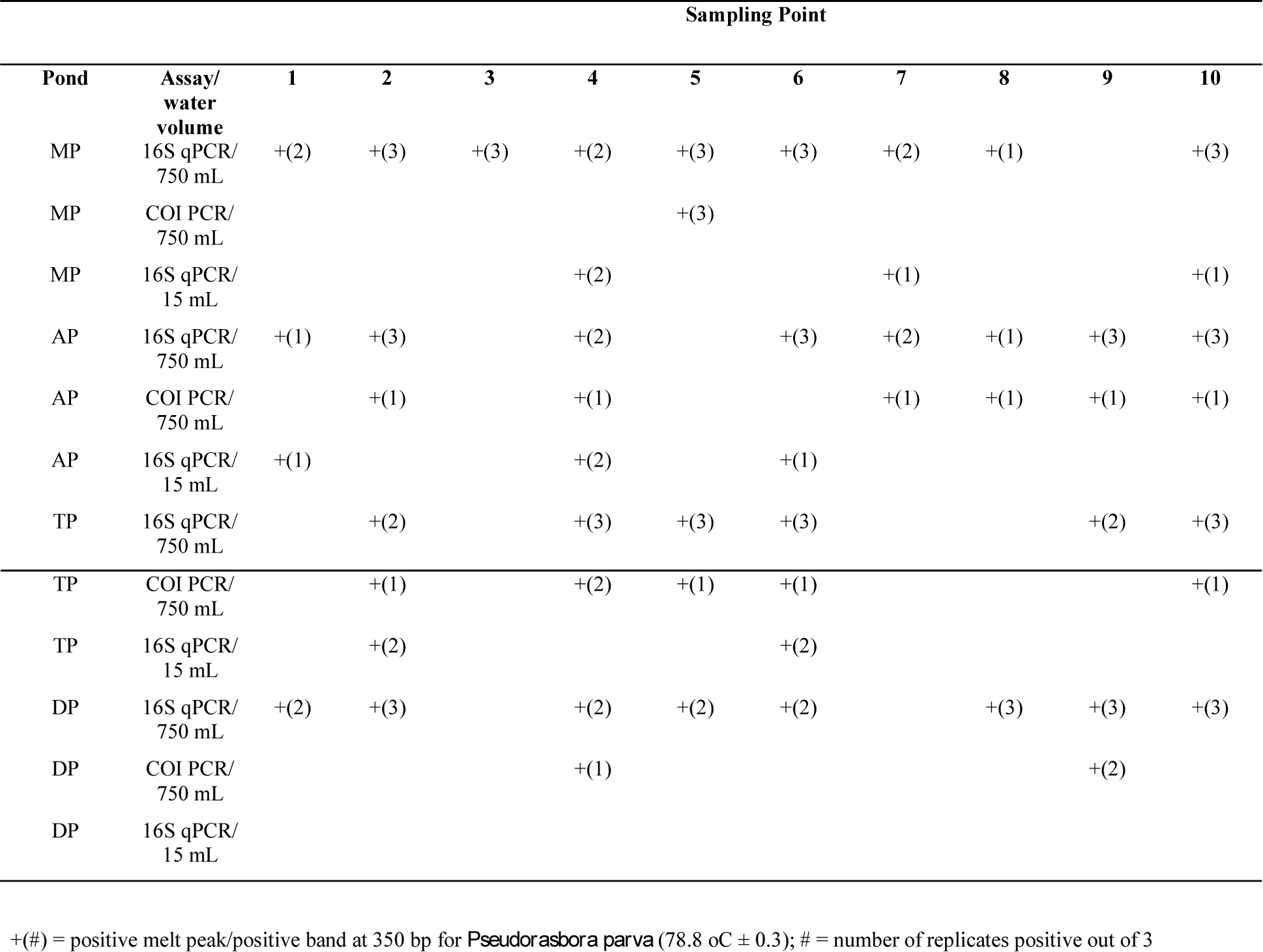
Positive amplifications for *Pseudorasbora parva* in filtered 750 mL water samples amplified with designed Pparva16S primers, 750 mL water samples amplified in end-point PCR with Davison *et al*. (2017) primers and positives amplifications in 15 mL water samples amplified with designed Pparva16S primers from all four ponds (Morolwg Pond, MP; Ashpits Pond, AP; Turbine Pond, TP and Dyfatty Pond, DP) at each sampling point (1 - 10) in optimised SsoFast™ EvaGreen^®^ qPCR assay. +(#) = positive melt peak/positive band at 350 bp for *Pseudorasbora parva* (78.8 oC ± 0.3); # = number of replicates positive out of 3

**Figure 2.**
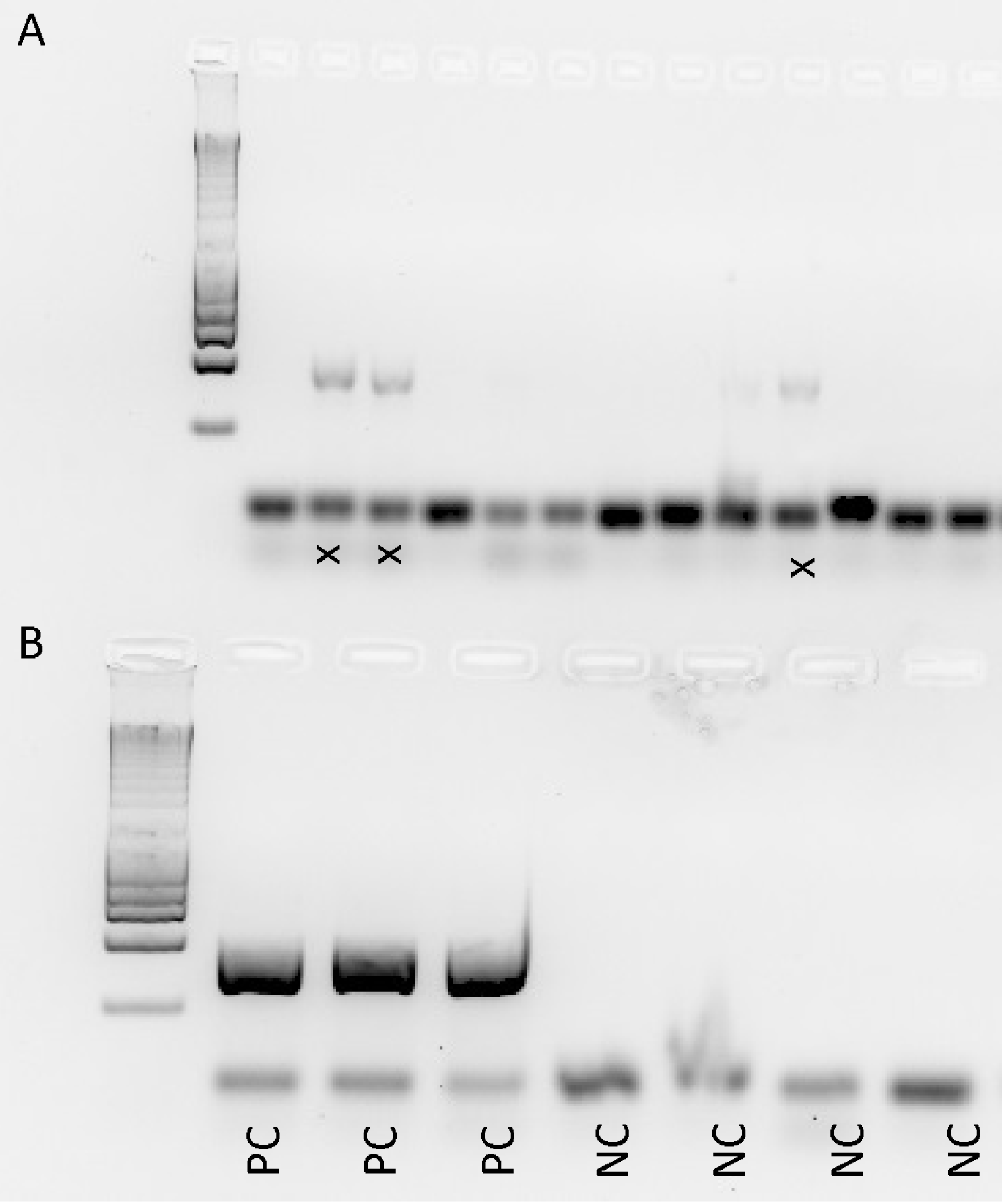
Agarose gel from end-point PCR using Davison et al., 2017 primers on field eDNA samples in triplicate for *Pseudorasbora parva.* A) Field samples (x) with positive samples displaying a band at 350 bp (200 bp ladder). B) Positive control tissue from topmouth gudgeon and negative controls.

Detection success varied significantly depending on eDNA assay (deviance = 31.32, df = 2, *P* <0.001) but not on pond identity (deviance = 3.36, df = 3, *P* =0.582l; Figure 3). Our novel qPCR 750 mL eDNA assay detected the presence of TMG (31/40 or 77.5%) in a significantly higher proportion of sites (*t* = 2.962, *P* = 0.016) than conventional PCR with the same water volume (14/40 or 35.0%) or qPCR with 15 mL of water (8/40 or 20.0%). The novel assay was 4.2 times more likely to detect topmouth gudgeon in an individual water sample than conventional end-point PCR, and 2.2 times more likely to detect its presence at a sampling site when multiple samples are collected (Table S5).

**Figure 3.**
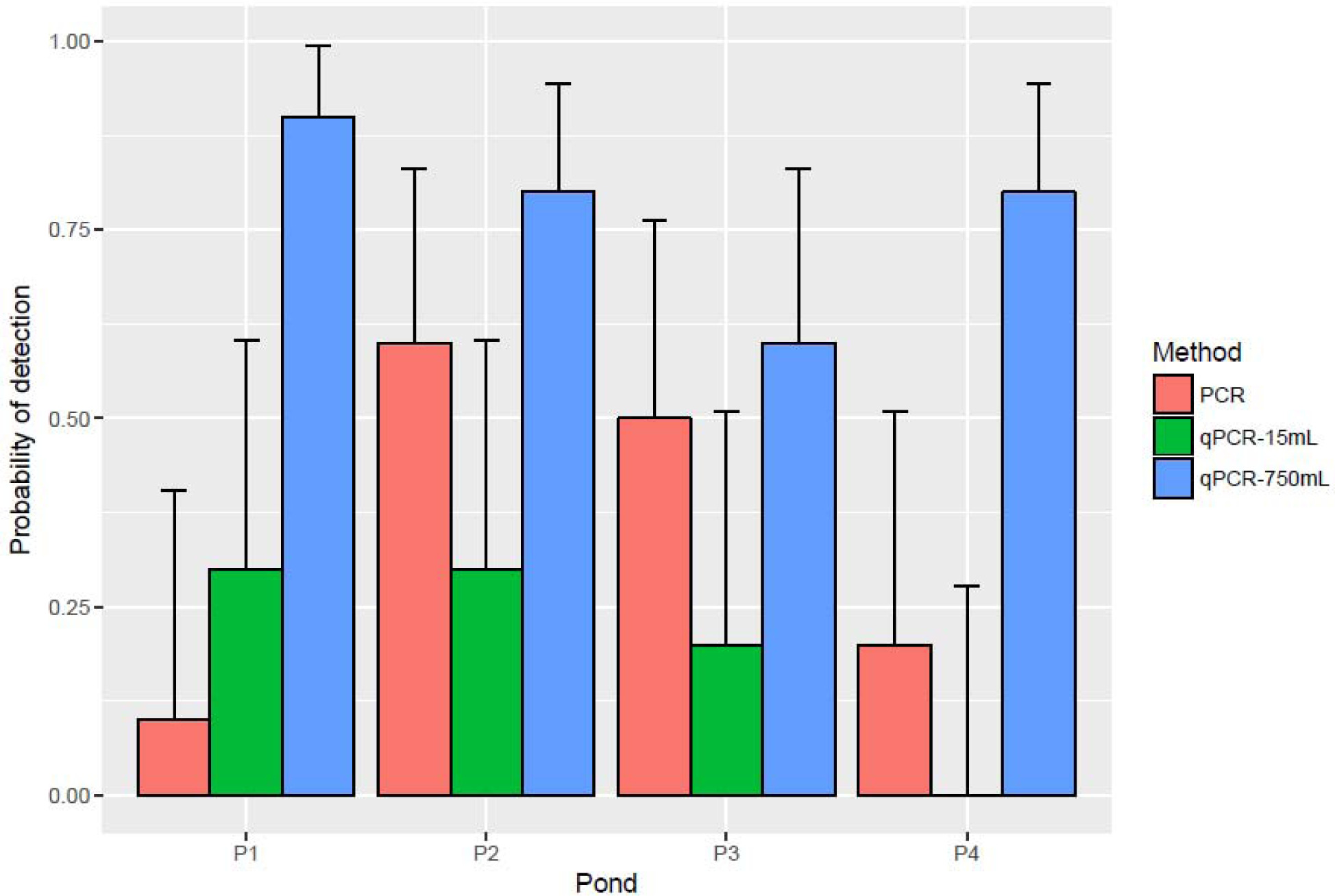
Probability of detection (± 95 binomial CI) of top mouth gudgeon (*Pseudorasbora parva*) at each pond (P1 - Morolwg Pond, 2 - Ashpits Pond; P3 - Turbine Pond; P4 - Dyfatty Pond) using different eDNA assays (end point-PCR on 750 mL of water, qPCR on 15 mL of water, and qPCR on 750 mL of water).

## Discussion

The application of our novel qPCR TMG assay detected the presence of TMG DNA at sites where the species was thought to have been eradicated or had not been detected by trapping. This serves to highlight the difficulties of inferring species absences from survey data (Jerde et al., 2011; Ficetola et al., 2015) and the superior sensitivity of qPCR-based eDNA methods over traditional approaches. No evidence of PCR inhibition was detected compared to previously published end-point PCR primers (Davison et al. 2017). The advantages of using an HRM-qPCR approach over end-point PCR include increased sensitivity and quantifiable levels of eDNA detection. In addition, the qPCR assay described here successfully amplified the target species in very small volumes of sample water (15 mL), albeit with lower sensitivity, which should greatly facilitate the collection of multiple replicated field samples, particularly in remote/inaccessible areas.

The use of eDNA to detect and monitor invasive species at low densities has numerous advantages over traditional techniques, namely higher sensitivity and ease of collection (Evans et al. 2017; Jones 2013; Tucker et al. 2016). For example, eDNA from European weather loach (*Misgurnus fossilis*) and redfin perch (*Perca fluviatilis*) was detected at sites where fishing had previously failed to find these species (Bylemans et al. 2016; Sigsgaard et al. 2015). TMG has a small size, and trapping can on occasions be unsuccessful, particularly during the early life stages (Jerde et al. 2011; Magnuson et al. 1994). The detection threshold for TMG in closed systems using traditional methods (electrofishing and traps) is approximately 0.5 fish per m^2^, which could suggest low TMG densities in both Ashpits and Dyfatty ponds (Britton et al. 2011b). The existence of false-negatives poses a particular problem for the management of AIS because they tend to occur at low population abundance, particularly during the early stages of invasion (Fitzpatrick et al. 2009). False negatives can also occur during the analysis of eDNA in the laboratory, due to poor primer design, low abundance of target DNA, and/or inadequate amplification (Ficetola et al. 2015). There are a number of benefits of using qPCR over traditional end-point PCR for the detection of target species using eDNA, such as greater sensitivity and quicker results (Nathan et al. 2014; Wood et al. 2013), and our results show that high resolution melt curve analysis (HRM) based on species-specific melt curve profiles (Héritier et al. 2017; Jaiswal et al. 2017) offers even greater sensitivity.

The volume of environmental samples collected can greatly influence the rate of detection of eDNA from a range of aquatic species (Pilliod et al. 2013; Rees et al. 2014). There is a trade-off between sampling effort (number of samples collected and processed) and probability of detection species (Rees et al., 2014). In our study, the probability of detection by qPCR was c. 4-6 times higher with a large water volume (750 mL) than with a smaller (15 mL) water volume, which is not unexpected considering the predicted low abundance of target DNA, the 50x reduction in water volume and the large size of the ponds sampled (Table 1, Table S5). Collecting smaller volumes of water with more replicates could offset the problem of sampling ponds and reservoirs without much additional effort (Goldberg et al. 2015; Rees et al. 2014) but our study shows that detection probabilities also depend on spatial scale, which will require careful planning.

We have demonstrated that eDNA screening can help detect invasive fish species, presumably occurring at low abundances. Low temperatures (<10 ^°^C), short trapping time (<4 hours), shallow deployment and below-threshold densities could explain the absence of adult TMG in the 2017 trapping, while size-selection biases can explain their absence during the 2016 surveys (Britton et al. 2011a). Post-eradication survey methods at the Millennium Coastal Park, including micromesh seine netting and trapping (plastic bottle and big mesh ‘minnow’ traps) is highly size-selective and it is possible that smaller colonising individuals (<20 mm) could have evaded nets and traps, resulting in false negatives and an incorrect indication of eradication success (Davies, Britton 2015). Even if eradication had been initially successful, it is possible that proximity to infected sites could have allowed fish to disperse through interconnecting streams (Britton et al., 2008; Copp *et al*., 2010; Pinder et al., 2005) and/or during flooding events, as reported for other AIS (Diez *et al.,* 2012; Rahel and Olden, 2008; Scott *et al.,* 2016). Inadvertent translocation of eggs and small larvae by local anglers across ponds is also possible as shown previously for this and other AIS (Britton et al. 2007; Johansson et al. 2018; Pinder et al. 2005).

In summary, routine monitoring with the assay developed here can provide valuable data regarding the expansion and dispersal of invasive TMG populations and, with further optimisation and calibration, could also be used to obtain relative estimates of species abundance. We have shown how the application of eDNA-qPCR methods can be used to monitor eradication programmes and help inform risk management strategies for TMG and other AIS under the Water Framework Directive.

## Acknowledgements

We would like to thank Gereint Mortimer (Afan Valley Angling Club) and Chris Lloyd for providing fish tissue samples for qPCR optimisation, Teja Muha for assistance in processing of DNA samples, Dr. Peter Jones for time and effort trapping topmouth gudgeon, and Emma Keenan (Natural Resources Wales) for assisting with eDNA sample collection and providing information on sampling sites. This research was funded by the Welsh Government and Higher Education Funding Council for Wales (HEFCW) through the Sêr Cymru National Research Network for Low Carbon Energy and Environment (NRN-LCEE; AQUAWALES cluster) and the EU Horizon 2020 AMBER project (www.amber.international) under EC grant agreement No 689682.

## Author contributions & competing interests

SC & CVR designed the study; CVR performed the analyses with advice from SC; CGL performed statistical analyses; MR carried out the field work and contributed to the writing; CVR, SC and CGL wrote the paper. Authors declare that they have no competing interests.

